# Sac1 links phosphoinositide turnover to cryptococcal virulence

**DOI:** 10.1101/2024.01.18.576303

**Authors:** Elizabeth A. Gaylord, Hau Lam Choy, Guohua Chen, Sydney L. Briner, Tamara L. Doering

**Affiliations:** Department of Molecular Microbiology, Washington University School of Medicine, St. Louis, MO, USA; L.E.K. Consulting, Boston, MA, USA; Pfizer, Inc., Chesterfield, MO, USA

**Keywords:** *Cryptococcus neoformans*, mycology, phosphoinositide, PI4P, phosphatase, Sac1, virulence

## Abstract

*Cryptococcus neoformans* is an environmentally-acquired fungal pathogen that causes over 140,000 deaths per year. Cryptococcal infection occurs when infectious particles are deposited into the lung, where they encounter host phagocytic cells. *C. neoformans* may be engulfed by these phagocytes, an important step of infection that leads to outcomes ranging from termination of infection to cryptococcal dissemination. To study this critical process, we screened approximately 4,700 cryptococcal gene deletion mutants for altered uptake, using primary mouse and human phagocytic cells. Among the hits of these two screens, we identified 93 mutants with perturbed uptake in both systems, as well as others with differences in uptake by only one cell type. We further screened the hits for changes in thickness of the capsule, a protective polysaccharide layer around the cell which is an important cryptococcal virulence factor. The combination of our three screens yielded 45 mutants, including one lacking the phosphatidylinositol-4-phosphate phosphatase Sac1. In this work, we implicate Sac1 in both host cell uptake and capsule production. We found that *sac1* mutants exhibit lipid trafficking defects, reductions in secretory system function, and changes in capsule size and composition. Many of these changes occur specifically in tissue culture media, highlighting the role of Sac1 phosphatase activity in responding to the stress of host-like conditions. Overall, these findings show how genome-scale screening can identify cellular factors that contribute to our understanding of cryptococcal biology and demonstrate the role of Sac1 in determining fungal virulence.

**IMPORTANCE:** *Cryptococcus neoformans* is a fungal pathogen with significant impact on global health. Cryptococcal cells inhaled from the environment are deposited into the lungs, where they first contact the human immune system. The interaction between *C. neoformans* and host cells is critical because this step of infection can determine whether the fungal cells die or proliferate within the human host. Despite the importance of this stage of infection, we have limited knowledge of cryptococcal factors that influence its outcome. In this study, we identify cryptococcal genes that affect uptake by both human and mouse cells. We also identify mutants with altered capsule, a protective coating that surrounds the cells to shield them from the host immune system. Finally, we characterize the role of one gene, *SAC1*, in these processes. Overall, this study contributes to our understanding of how *C. neoformans* interacts with and protects itself from host cells.

## INTRODUCTION

*Cryptococcus neoformans* is a fungal pathogen that causes significant mortality in immunocompromised individuals, particularly those with HIV/AIDS (1). Despite recent advances in healthcare access and treatments, cryptococcal meningitis still leads to 19% of AIDS-related deaths (2). The interaction between *C. neoformans* and host phagocytes begins when environmentally-acquired fungal cells are deposited in the lung. Subsequent phagocytosis may lead to fungal cell death and termination of the infection. However, in some cases cryptococcal cells can withstand the pressures exerted by phagocytes to survive and proliferate within phagolysosomes (3–6). In this scenario, engulfment of the yeast likely enhances cryptococcal dissemination from the lung to other sites (7). Due to the pivotal role of this initial phase of cryptococcal pathogenesis, extensive efforts have been devoted to studying the interactions of *C. neoformans* with host phagocytic cells.

*C. neoformans* produces multiple factors which modulate its interactions with host cells and thereby provide protection from phagocytosis. These include the cell wall and surrounding polysaccharide capsule, complex glycan-based structures which change in size and composition in response to environmental conditions. Strains with perturbations of these structures often exhibit increased uptake by phagocytic cells (8, 9), demonstrating the importance of capsule for protecting against phagocytosis.

Various models have been used to interrogate cryptococcal-host interactions. These include invertebrate and vertebrate animals, amoeba, and primary and immortalized cells (3, 10–16). Genetic screens based on libraries of mutant host or fungal cells have identified gene products that modulate interactions of cryptococci with specific cell lines. In one example, Chun and colleagues subjected cryptococcal cells to insertional mutagenesis and identified mutations that affected their uptake by RAW264.7 macrophages (17). Srikanta et al. used RNA interference and high-throughput imaging to find host regulatory genes which govern the interactions of *C. neoformans* with THP-1 cells and discovered 25 fungal-specific regulators of phagocytosis (18). By applying high-throughput imaging to a *C. neoformans* gene deletion collection (19), Santiago-Tirado et al. identified 56 fungal genes that affect uptake by THP-1 cells (9). A common strength of these studies is the genetic tractability and accessibility of the immortalized cell lines used. However, such lines lack some features of primary host cells (20–22), which may limit these approaches. Screens may also be constrained by the availability and size of *Cryptococcus* mutant collections. For example, the last study cited screened a library of approximately 1,200 single gene deletion mutants (9), whereas much larger libraries are currently available, increasing our power to identify novel phenotypes.

This report incorporates several important departures from previous screens designed to elucidate *C. neoformans*-host interactions. First, we screened a collection of 4,692 single gene deletion mutants, representing over eighty percent of the non-essential genes in the cryptococcal genome. Second, we applied a flexible medium-throughput imaging method that avoids pitfalls of earlier studies, such as difficulty distinguishing adherent versus internalized yeast cells (23). Third, we screened using primary cells from both mouse (murine bone marrow-derived macrophages; BMDM) and human (monocyte-derived macrophages; HMDM), to take advantage of a model system while also examining the system of greatest clinical interest. Finally, we reasoned that many of our hits would have altered capsules, so we further tested mutants that demonstrated altered phagocytosis for changes in capsule thickness. Overall, we identified 45 deletion strains which exhibited altered uptake in both systems as well as altered capsule size.

While it is known that capsule material traffics at least in part through the secretory pathway (24), few proteins involved in this process have been characterized. To address this gap, we focused on mutants that might have aberrant capsule trafficking. One critical process in the regulation of secretion is the subcellular distribution of phosphoinositides (PI). Various cellular PI species are defined by the patterns of phosphorylation on their inositol head groups. These species are maintained at distinct cellular compartments by kinases and phosphatases, which specifically and reversibly phosphorylate and dephosphorylate inositol (25). In this way, PIs help define membrane identity and regulate membrane-protein interactions (25), thus playing important roles in secretory trafficking (26, 27). We therefore were particularly interested to note that one strain, which emerged as a hit in all three of our screens, lacks the gene *SAC1* (*CKF44_06080*). Sac1 is a phosphatase that is highly conserved in eukaryotes and acts in phosphoinositide metabolism (28). Our observation that cells lacking Sac1 are hypocapsular thus suggested that modulation of phosphoinositide turnover in the secretory pathway influences capsule synthesis. Below we use biochemical, imaging, and molecular engineering approaches to establish the mechanism of this interaction, linking alterations in lipid homeostasis to the production of a critical cryptococcal virulence factor.

## RESULTS

### Fungal genes that influence interactions with host cells

To identify fungal gene products that affect interactions with host cells, we screened 4,692 *C. neoformans* single gene deletion mutants (19). In our assay, fungal cells were stained with Lucifer Yellow, opsonized with serum, and then incubated with primary phagocytes to allow engulfment. The samples were washed to remove unassociated cells and then stained with membrane-impermeant Calcofluor White to label externally-associated fungi and propidium iodide to label phagocytic cell nuclei. We assessed fungal engulfment as the phagocytic index, or number of internalized fungi (stained with Lucifer Yellow but not Calcofluor White) per host cell (identified by nuclear staining) (23). We define the uptake score as the ratio of a mutant’s phagocytic index to that of wild-type *C. neoformans* (KN99).

We first applied our assay to screen the interactions of *C. neoformans* with murine bone marrow-derived macrophages (BMDM), normalizing to fungal cell number to account for differences in mutant growth rates (Figure 1A). Because these deletion library plates do not include WT control cells, we initially compared strain uptake scores to the median for each plate and used permissive cutoffs to call roughly 10% of the strains as hits (Table S1A). We then subjected the 458 strains that we identified with increased or decreased uptake (Table S1B) to a second round of BMDM screening (Table S1C), this time comparing them to in-plate wild-type controls and using more stringent cutoffs to select hits (Figure 1B; Table S1D). In parallel, we screened them using human monocyte-derived macrophages (HMDM) in place of BMDM (Figure 1C; Tables S1E and S1F).

**FIgure 1.**
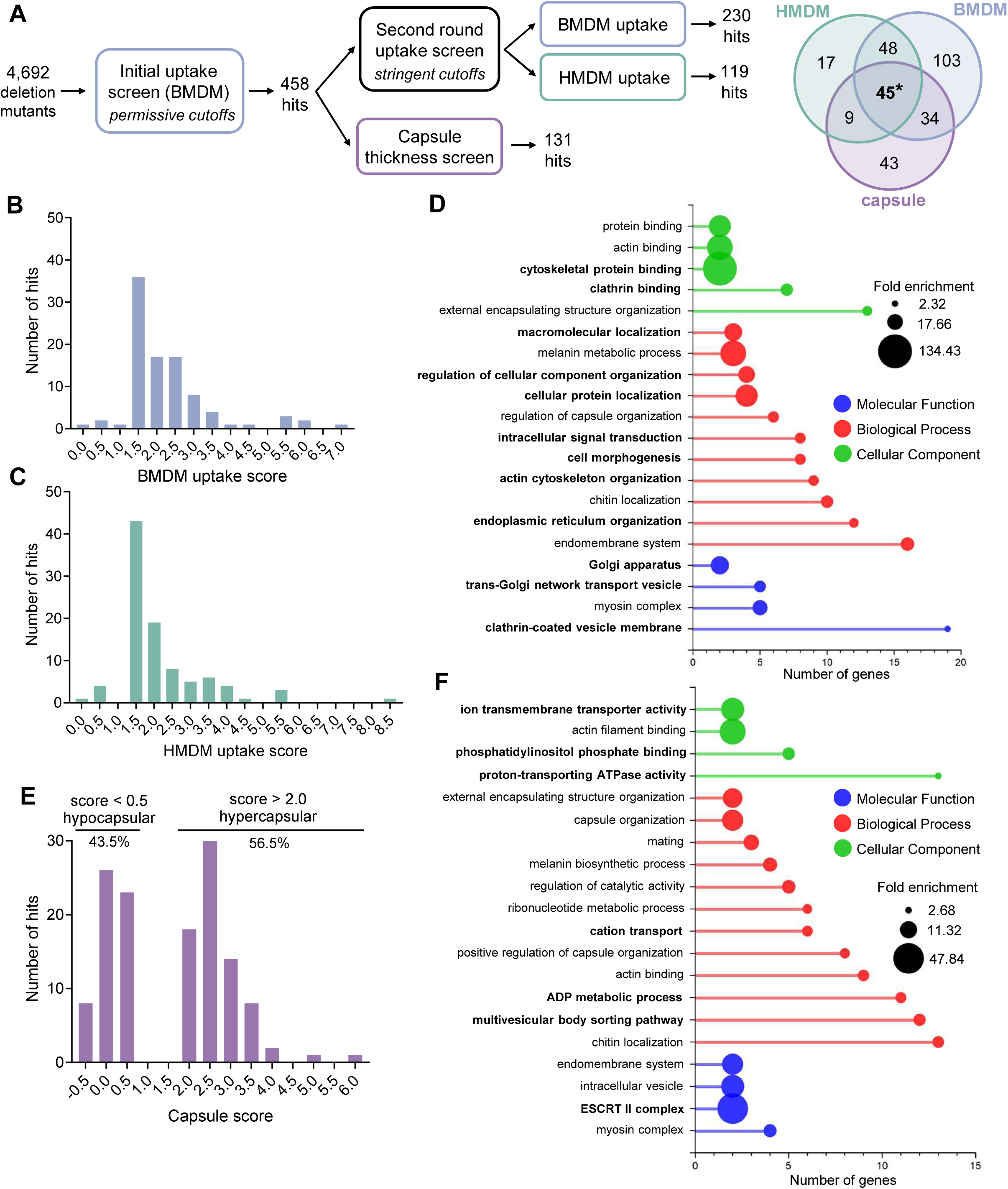
Screen results. **(A)** Left, schematic of screening workflow, and right, overlap of screen hits. Asterisk indicates *SAC1* in diagram at right. **(B and C)** Uptake scores (as defined in the text) for the 93 mutants identified as hits when screened against **(B)** BMDM and **(C)** HMDM. The value under each bar represents the center of the bin and 1.0 is the WT value. **(D)** Selected terms that are enriched in the 93 dual hits from the uptake screen. X-axis, number of hit gene IDs which fall within each gory. **(E)** Capsule score (capsule thickness relative to that of WT under inducing conditions) for 131 mutants identified as hits e capsule screen, plotted as in **(B)** and **(C)**. **(F)** Selected GO terms that are enriched in hits from the capsule screen, plotted as in **(D)**. For panels D and F, bold text denotes categories uniquely enriched for each screen; note that not all categories are listed.

Our second round of screening narrowed the initial hits to 93 mutants that significantly influenced host interactions in both BMDM and HMDM cells (Figure 1A, overlap of red and blue circles; Table S1G). When we applied Gene Ontology (GO) analysis to the corresponding genes, we found that genes encoding cellular factors related to cytoskeletal interactions and intracellular protein localization and transport, among other categories, were particularly enriched relative to their abundance in the genome (Figure 1D; see Discussion).

### Mutants with altered host interactions and capsule size

We were particularly interested in gene products that might act in capsule production, because of the many outstanding questions that remain about this process (29). As the outermost layer of the cell, the capsule is fundamental in mediating cryptococcal interactions (7). Therefore, we hypothesized that our uptake screen hits would be enriched for mutants with altered capsule. To test this idea, we adapted an automated, fluorescence-based capsule quantification assay (30) to accommodate medium-throughput screening. Of the 458 first round hits from the uptake screen, 131 indeed showed capsule changes (Figure 1A), with 57 strains identified as hypocapsular and 74 as hypercapsular (Figure 1E; Tables S2A, 2B). Validating the screen, this group contained multiple previously-identified capsule mutants, including 16 that were hypocapsular and 3 that were hypercapsular. GO terms for the hits were particularly enriched for roles in capsule organization, as we expected, and for processes related to vesicular traffic, some of which were also enriched in our uptake screen hits. Uniquely enriched for the capsule screen hits were ion transport and phosphoinositide binding (Figure 1F).

We identified 45 mutants which were hits in all three of our screens (Table S2C). The combination of perturbed interactions with phagocytes and altered capsule thickness of these strains suggested that the deleted genes would be important for virulence. We further speculated that the uptake phenotypes were related to surface changes in the cells, which might be mechanistically elucidated by analysis of the deleted genes. Of the 45 triple hits, 18 had been previously characterized. When we used homology to predict the subcellular localization of the remaining gene products, the largest group (8 of 27) was predicted to reside in the secretory pathway. We suspected these genes might influence secretion of capsule material.

Among the eight mutants likely impaired in secretion, we particularly noticed one strain that showed increased phagocytosis by both BMDM and HMDM and a 61% reduction in capsule thickness compared to WT cells (Figure 2A). Its missing sequence, *CKF44_06080*, encodes a protein with 38% amino acid identity to the phosphatidylinositol phosphatase Sac1 of *Saccharomyces cerevisiae,* including 85% identity across a 20-amino acid region that encodes a conserved CX_5_R(RT) motif necessary for catalytic function (31). Interestingly, despite this similarity, ectopic overexpression of *S. cerevisiae* Sac1 did not restore the capsule defects of *sac1*Δ cells (Figure S1).

**Figure 2.**
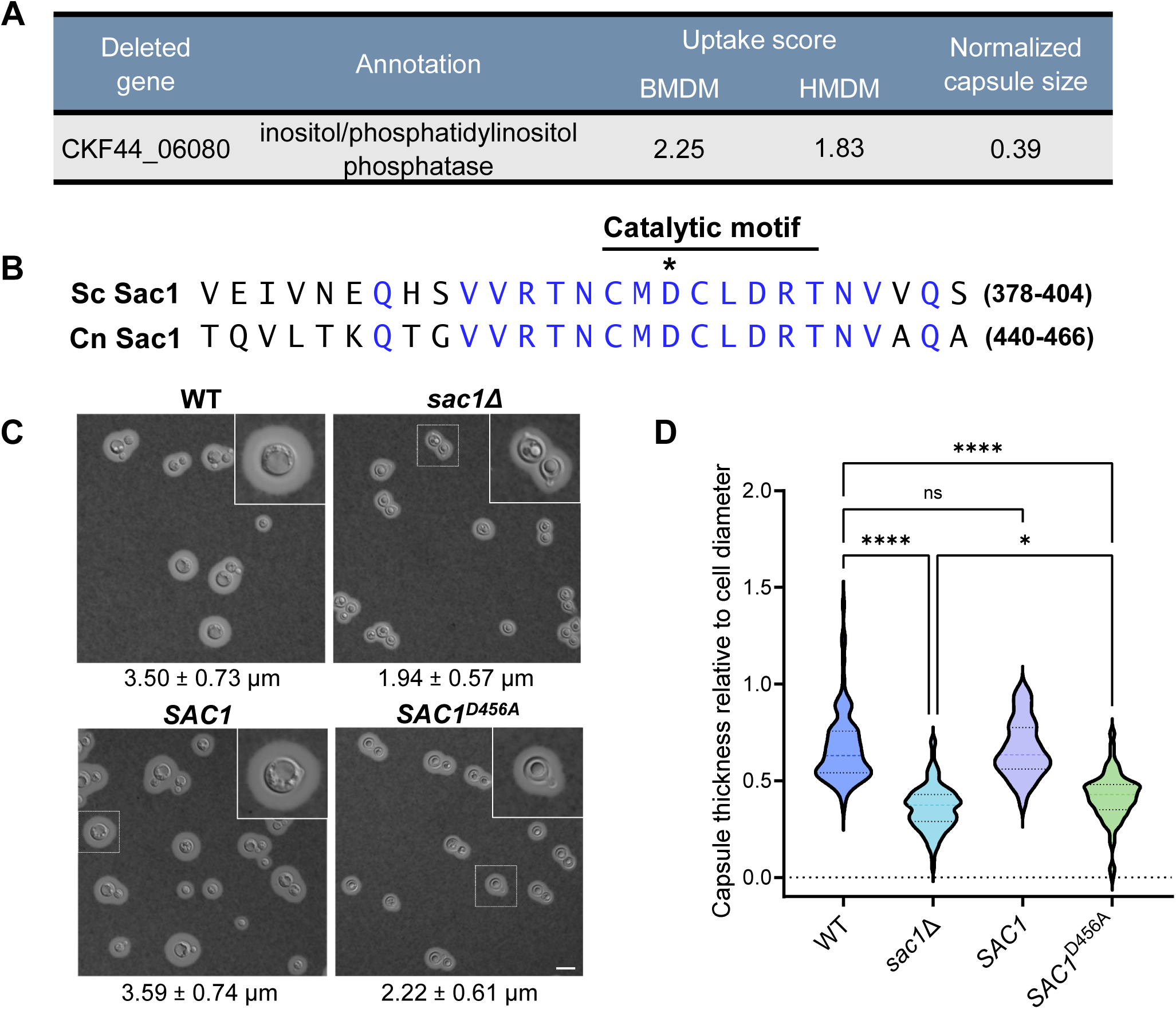
Cells lacking active Sac1 show increased uptake by phagocytes and smaller capsules. **(A)** Uptake and capsule screen results for cells lacking CKF44_06080. **(B)** Alignment of *C. neoformans* and *S. cerevisiae* sequences that include the Sac1 catalytic motif. Blue, identical residues; asterisk, residue mutated to alanine in the *SAC1^D456A^* strain. **(C)** India ink negative stain of the indicated strains grown for 24 hours in DMEM at 37°C with 5% CO_2_. Capsule thickness in μm (mean ± SD) is noted below each panel. All primary images are at the same magnification; scale bar, 10 μm. For this and other microscopy figures, any region demarcated by a small square is enlarged in the inset at upper right. **(D)** Distribution of normalized capsule thicknesses for the indicated strains. At least 96 cells were measured for each strain as described in the Methods. *, p<0.0332; ****, p<0.0001 by 2-way ANOVA with Tukey’s multiple comparison test; ns, not significant.

The phenotypes of *sac1*Δ cells, together with the unique biology of the capsule and its importance as a virulence factor, led us to further investigate the role of Sac1 in *C. neoformans*. To pursue these studies, we first engineered a new Sac1 deletion strain (*sac1*Δ) and complemented the mutant at the endogenous site (*SAC1*). We also identified the conserved Sac1 catalytic motif by homology to *S. cerevisiae* Sac1 and inactivated it by replacing an aspartic acid residue with alanine (*SAC1^456A^*, Figure 2B and S2) (32, 33). Using these strains, we confirmed that loss of Sac1 led to thinner capsules, a phenotype which was maintained after normalization to the smaller diameters of the *sac1*Δ cells (Figure 2C, D). These traits were phenocopied by the inactivated strain, showing that the phosphatase function of Sac1 is essential for capsule production, while complementation restored WT phenotypes (Figure 2C, D). While imaging, we also observed striking round structures in the *sac1* mutant cells (Figure 2C, right panels). These were distinguished from typical organelles, like the vacuoles visible in WT and *SAC1* cells, by their circular shape, defined edge, and striking refractility and only occurred in *sac1* mutants grown in host-like conditions (here defined as DMEM, 37°C, 5% CO_2_). These structures will be further discussed below.

### Sac1 is required for cryptococcal virulence

Variations in both uptake and capsule size are associated with changes in pathogenicity (9, 34–40), so we suspected that *sac*1 mutants would have impaired virulence. To test this, we infected mice intranasally and measured organ burden at 14 days post-infection. Notably, while *sac1*Δ cells were recovered from the lungs, they remained close to the level of the original inoculum. In sharp contrast, the lung burden of WT and *SAC1* infected animals increased roughly 2000-fold in this period (Figure 3A). Cells were recovered from the brain in only a minority of mutant infections, although all WT-infected animals had detectable brain burden by this time point (Figure 3B). We observed similar results in a time course study, which demonstrated gradual accumulation of cryptococci in the lung up to 18 days post-infection (Figure S3A) and impaired dissemination to the brain (Figure S3B). These results motivated us to define the mechanism(s) by which Sac1 influences virulence.

**Figure 3.**
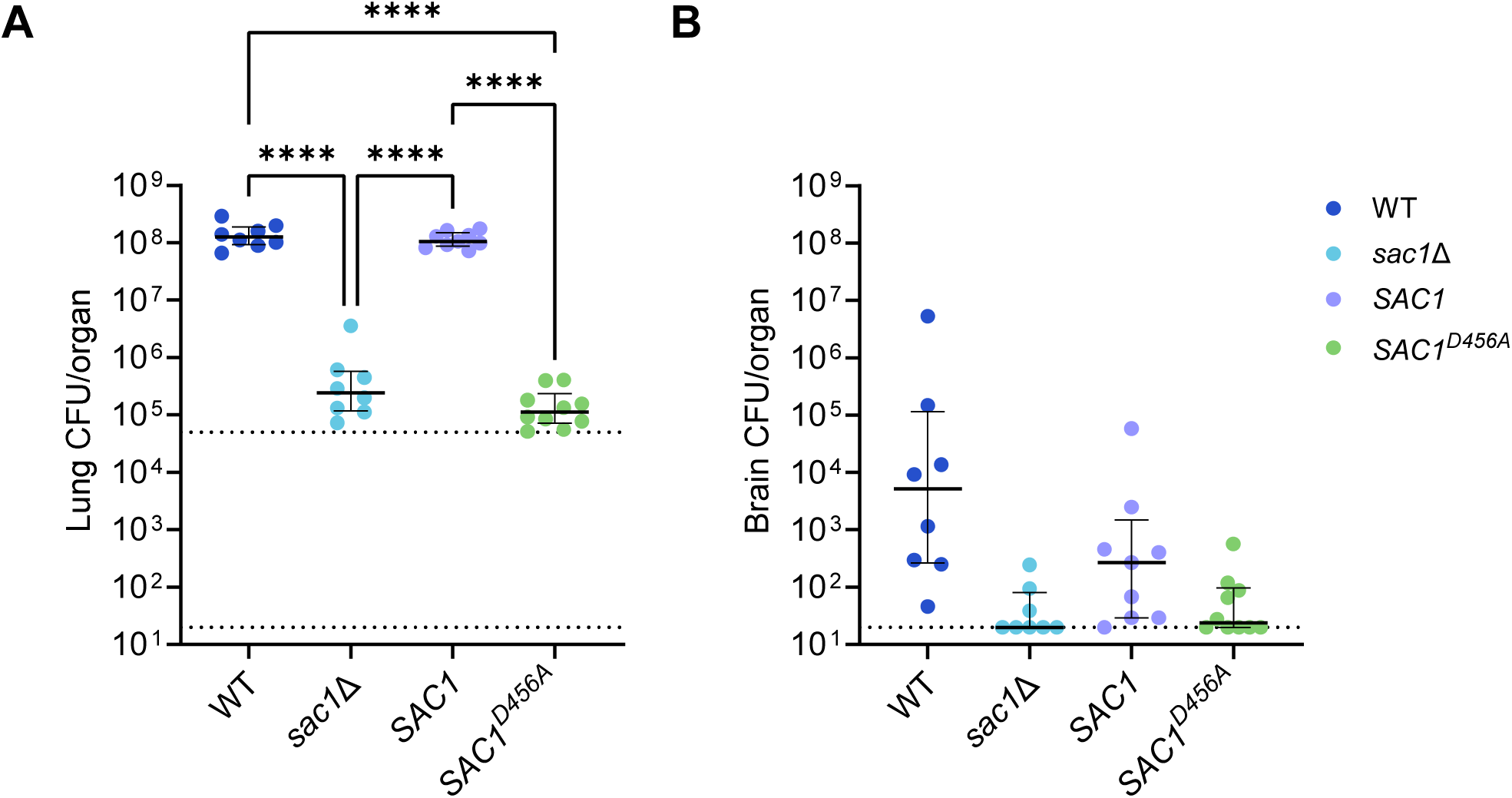
Sac1 is important for virulence in a mouse infection model. **(A)** Lung and **(B)** burden of C57BL/6 mice (groups of 8-10) sacrificed 14 days after intranasal infection with 5×10^4^ fungal cells (upper dotted line). Plots show median (bold black line) with interquartile range. Lower dotted line, limit of detection. Each symbol represents one mouse. ****, p<0.0001; comparison with no stars shown are not significant. Significance was calculated using 2-way ANOVA with Tukey’s multiple comparison test.

### Sac1 controls substrate distribution and sensitivity to stressors

Sac1 has not been characterized in *C. neoformans.* To test its influence on turnover of intracellular PI4P, we engineered strains to express an mNeonGreen-tagged peptide that binds PI4P (FAPP1) and has been used to assess its localization (41). We then examined the abundance and localization of this fluorescent peptide (Figure 4A). Overall, the FAPP1 signal was brighter in *sac1*Δ cells than in WT, a trend supported by flow cytometry (Figure S4). This is consistent with a role for Sac1 as a PI4P phosphatase, as loss of this protein would lead to an increase in its substrate. We also noted distinctive bright perinuclear rings in the *sac1*Δ cells (especially in rich medium), which were more diffuse in WT cells. These correspond to an endoplasmic reticulum distribution (42) and suggest aberrant accumulation of PI4P in the early secretory pathway. Finally, the mutant strain exhibited intense puncta, absent from WT cells, which are consistent with PI4P accumulation in the yeast Golgi (43).

**Figure 4.**
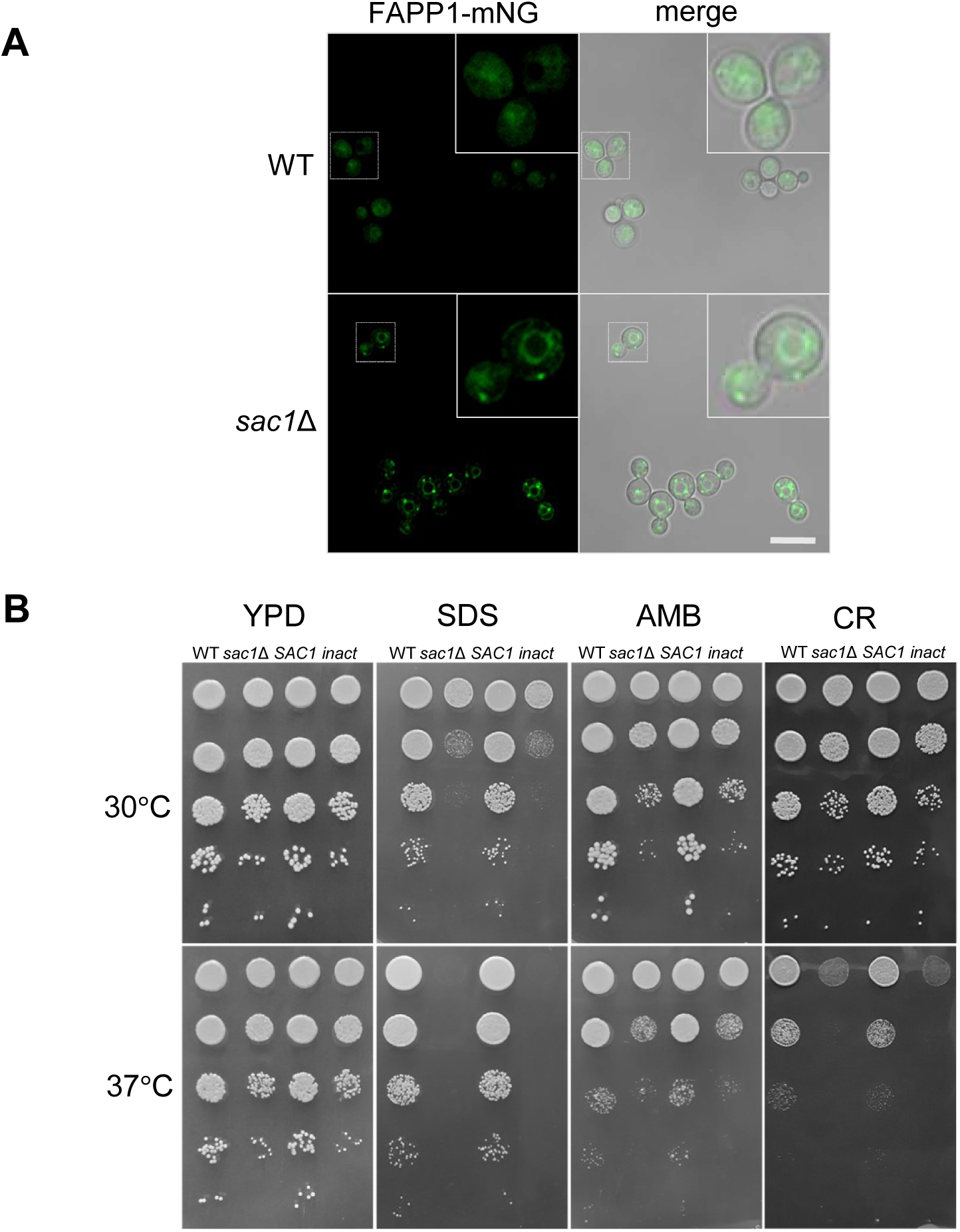
Sac1 regulates substrate distribution and stress sensitivity. **(A)** FAPP1-mNeonGreen (mNG) localization in cells grown in YPD at 30°C. All primary images are at the same magnification; scale bar, 10 μm. **(B)** Serial 10-fold dilutions of the indicated strains plated on YPD alone or containing 0.01% SDS (SDS), 0.05% Congo Red (CR), or 2 μg/mL Amphotericin B (AMB), incubated at 30 and 37°C. *inact* = *SAC1^D456A^*.

Secretion is a vital cellular process, so its perturbation could have multiple deleterious effects on the cell, including effects on the plasma membrane and cell wall. To examine this, we tested strain growth in various stressful conditions. Although both *sac1* mutants grow slightly slower than WT or complemented cells even on rich medium, they showed much worse growth on media containing membrane stressors (the detergent SDS and the sterol-binding antifungal amphotericin B) or the cell wall stressor Congo Red (Figure 4B). These defects were exacerbated when cells were grown at 37°C. In all of our assays, the complemented strain phenocopied WT and the inactivated construct phenocopied the deletion mutant.

### Sac1 regulates cellular lipids during the response to host-like conditions

We sought to explain the capsule defect and stress sensitivities of *sac1* mutants based on the cellular consequences of perturbed PI4P turnover. In other eukaryotes, PI4P is important for establishing the lipid composition of secretory vesicles (44) and its turnover is critical for the progression of post-Golgi secretory traffic (45). We therefore examined classes of molecules, including lipids, proteins, and capsule polysaccharides, that are directly or indirectly influenced by this step of secretion. As noted above, we had observed formation of large internal structures in *sac1* mutant cells grown under host-like conditions (Figure 2C), which were absent from WT and complemented cells. Because of their shape and refractile appearance in light micrographs, we suspected they were at least in part composed of lipid. Consistent with this suggestion, they appeared electron-lucent by transmission electron microscopy (Figure 5A, asterisks). We hypothesized that these structures result from aberrant lipid handling within *sac1* mutant cells that is induced by the stress of host-like conditions, since they are absent from mutant cells grown in rich medium (Figure 5A). Based on this idea, and our results below, we termed them Sac1 lipid-accumulating bodies (SLBs).

**Figure 5.**
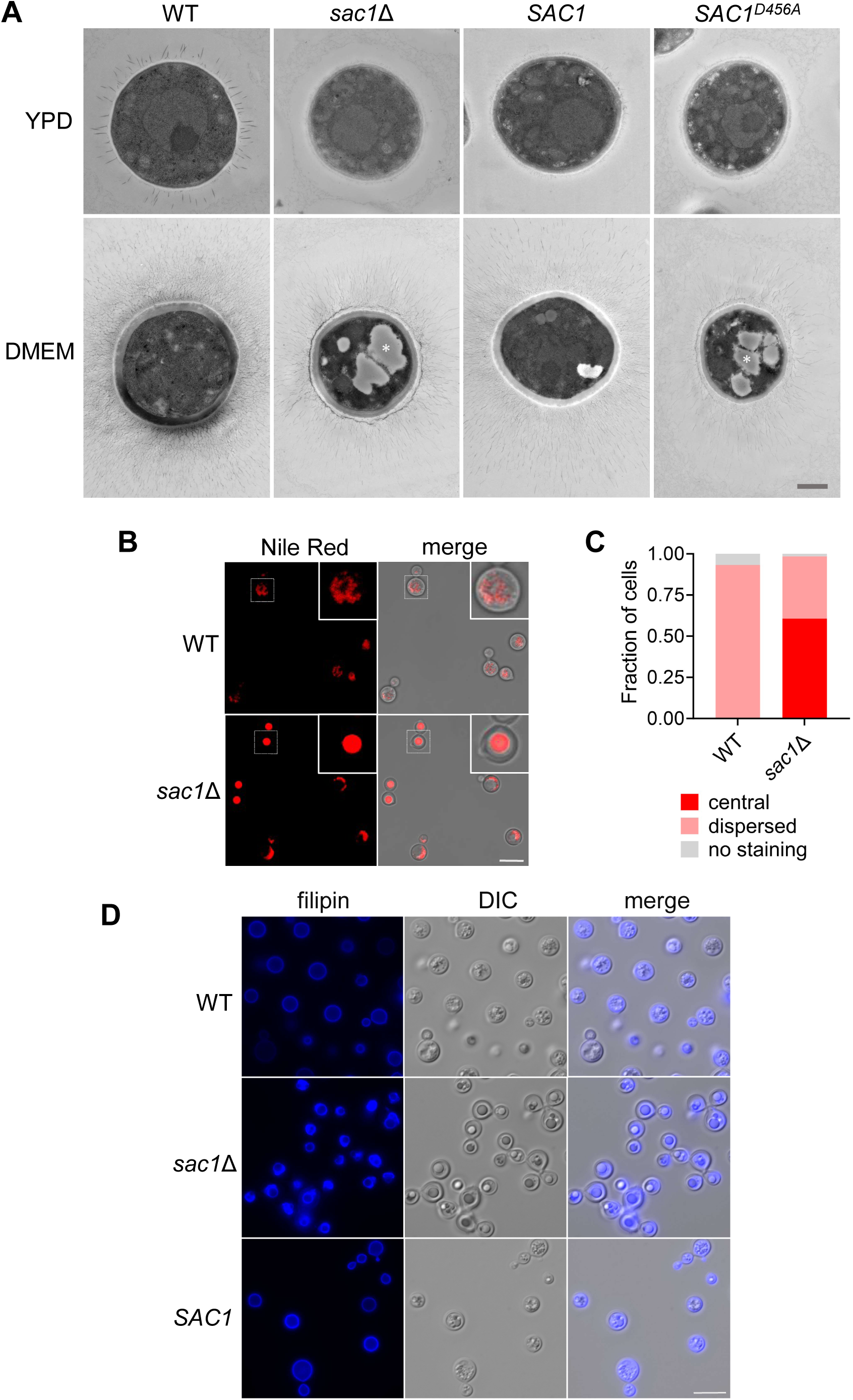
Sac1 is essential for proper lipid trafficking in host-like conditions. **(A)** Transmission electron micrographs of cells grown in the indicated medium. White asterisks, example lipid droplet structures, likely distorted during sample prep; scale bar, 1 μm. **(B)** Nile Red stain of the indicated strains with **(C)** quantification of the observed staining pattern. **(D)** Filipin staining of cells grown in DMEM. YPD grown cells in Panel A were grown overnight at 30 °C, all other cells were grown for 24 h in DMEM at 37°C with 5% CO_2_. Scale bar, 10 μm. All images within each experiment are shown at the same magnification.

SLBs resemble previously-characterized ”supersized” lipid droplets identified in *S. cerevisiae* (46). Lipid droplets are neutral lipid-containing structures that represent essential cellular energy reserves; these increase in abundance when cryptococcal cells are grown in host-like conditions (47). When we stained for neutral lipids using Nile Red, WT cryptococcal cells showed dispersed punctate signal, while *sac1Δ* cells showed concentrated central staining, frequently colocalized with SLBs (Figure 5B,C) and thus supporting the presence of neutral lipids within them.

We recently demonstrated that mislocalization of the important fungal lipid ergosterol leads to accumulation of lipid droplets (47). We had also noted that *sac1* mutants were more sensitive than WT cells to Amphotericin B, which binds ergosterol (Figure 4B). Consequently, we wondered whether ergosterol localization in *sac1*Δ cells was perturbed. When we stained WT cells grown in host-like conditions with filipin, which binds non-esterified sterols, we saw the typical ring-staining expected for a component of the plasma membrane (Figure 5D). By contrast, mutant cells exhibited increased internal staining, particularly bordering the SLBs (see Discussion). When we tested filipin and Nile Red together, both stains localized to SLBs in the *sac1* mutants, with a ring of filipin surrounding the Nile Red stain (Figure S5). In contrast, WT cells showed peripheral filipin staining and Nile Red staining dispersed throughout the cell.

In testing cell growth, we fortuitously discovered that supplementation of host-like conditions (DMEM, 37°C, 5% CO_2_) with various exogenous fatty acids partially restored the growth of *sac1*Δ cells (Figure 6 and Supplemental Figure S6); this effect was dose-dependent (shown for myristate in Figure 6A). This supplementation also prevented the formation of SLBs (Figure 6B, bottom right panel). When we visualized capsule in myristate-supplemented cells, we made several observations. First, the addition of the BSA vehicle alone led to a slight capsule size increase, of similar magnitude, in both cell types. Second, myristate supplementation yielded significant capsule reduction in both strains, which was more dramatic in the mutant (Figure 6B). Lipid supplementation thus restores growth and prevents SLAB formation in *sac1*Δ cells but does not rescue capsule size (see Discussion).

**Figure 6.**
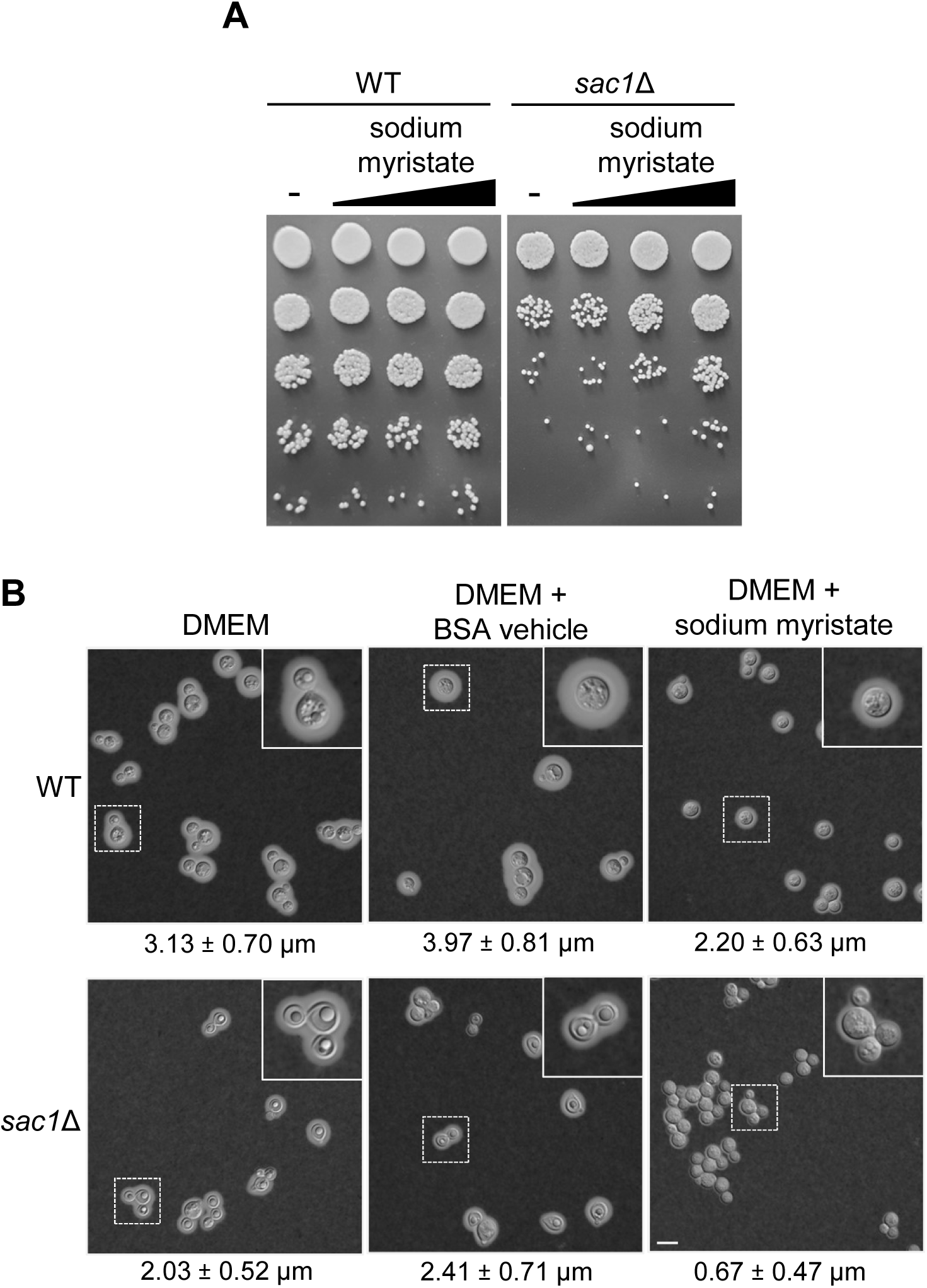
Supplementation with exogenous fatty acid partially restores *sac1*Δ growth and morphology but not capsule biosynthesis. **(A)** Serial 10-fold dilutions of WT and *sac1*Δ cells grown for 24 hours (at 37°C with 5% CO_2_) in DMEM supplemented with BSA alone (-), or with 8, 40, or 200 μM myristic acid conjugated to BSA (increasing amounts indicated by the triangle). **(B)** India ink negative stain of WT and *sac1*Δ cells grown in DMEM at 37°C with 5% CO_2_, supplemented as indicated with BSA alone or 200 μM myristic acid conjugated to BSA. Capsule thickness in μm (mean ± SD) is noted below each panel. All primary images are to the same magnification; scale bar, 10 μm.

### Sac1 impacts protein secretion and localization

Proteins are critical secretory cargo, so it is likely that accumulation of PI4P in the secretory pathway would affect their transport and secretion. To assess this, we measured protein secretion directly, using inducible acid phosphatase production as a proxy for general protein secretion (24). Upon entry into phosphate-depleted conditions, WT, *SAC1*, and both *sac1* mutants began producing detectable levels of acid phosphatase within one hour. While the general patterns of secretion were similar, the mutants produced less acid phosphatase on a per cell basis throughout the time course (Figure 7A), suggesting an overall reduction in protein secretion.

**Figure 7.**
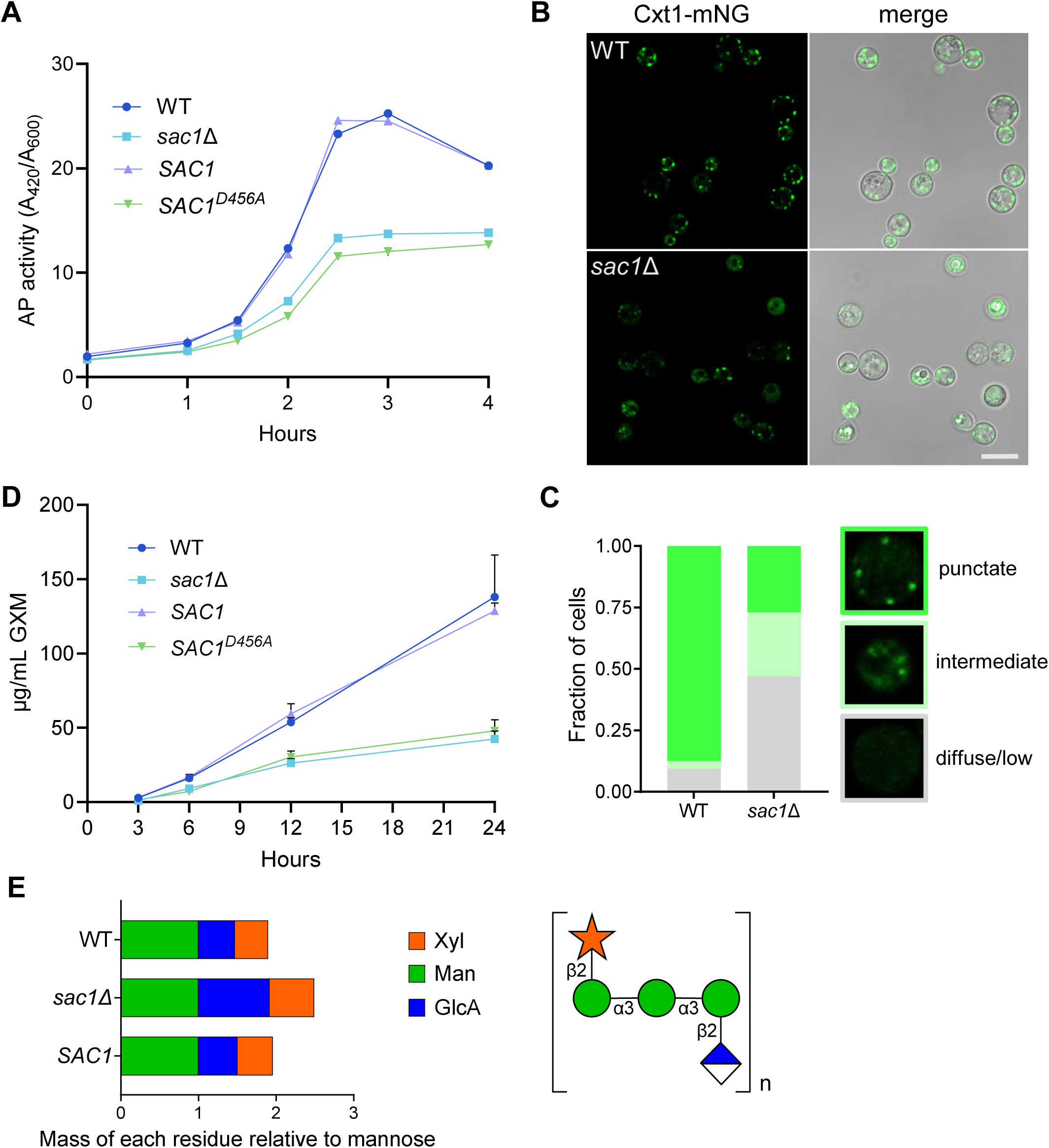
Sac1 controls the secretory pathway to modulate capsule size and composition. **(A)** Secreted acid phosphatase activity, normalized to cell number, over time after a shift to phosphate-depleted medium. **(B)** Confocal imaging of Cxt1-mNG in the indicated strain backgrounds with **(C)** quantification of observed staining patterns. Punctate: 3 or more puncta present; Intermediate: puncta and diffuse signal; Diffuse/low: diffuse signal or <3 puncta. Images are to the same scale; scale bar, 10 μm. **(D)** Shed GXM quantified by ELISA. For both **(A)** and **(D)**, mean ± SD for one representative experiment of three biological replicate studies is shown. **(E)** Left, GXM composition; right, subunit structure of GXM.

Perturbations in PI4P levels may affect not only proteins secreted outside the cell, but also those resident in the secretory system, including multiple capsule biosynthetic proteins in the Golgi apparatus. We hypothesized that the loss of PI4P turnover could result in failure to correctly localize these enzymes and thereby explain the altered capsule of *sac1* mutants. To investigate this question, we localized the best-characterized capsule biosynthetic enzyme, Cxt1 (48), in *sac1*Δ and WT cells grown in host-like conditions. WT cells displayed clear puncta of Cxt1-mNeonGreen, consistent with the expected Golgi localization of this protein. In contrast, staining in *sac1*Δ cells was heterogenous, with various cell populations displaying punctate, diffuse, or little to no fluorescence (Figure 7B, C).

### Sac1 affects capsule secretion and composition

Capsule glycans constitute a class of secretory cargo which is unique to *C. neoformans*. The predominant capsule polysaccharide, glucuronoxylomannan (GXM), traverses the secretory pathway prior to extracellular release (24). We hypothesized that GXM secretion, like protein secretion, is reduced in *sac1*Δ cells, causing their thin capsules. To test this, we quantified GXM secretion at intervals after transition to host-like conditions. Importantly, WT and *sac1* mutants began to produce detectable levels of GXM concurrently, showing that the mutants respond normally to the environmental signals that trigger capsule production (Figure 7D). However, the overall amount of GXM they secreted was markedly reduced.

Reduced GXM secretion could certainly explain the small capsules of *sac1* mutant cells. Another reason for altered capsule could be mislocalization of glycosyltransferases (such as we observed with Cxt1), since compositional differences in the polysaccharide also affect capsule size and architecture (8). To investigate this, we analyzed the composition of GXM purified from WT, *sac1*Δ, and *SAC1* cells. Compared to the WT and *SAC1* controls, *sac1*Δ GXM showed a substantial (∼2-fold) increase in the relative proportion of glucuronic acid, as well as a modest increase in xylose (Figure 7E). These results were supported by linkage analysis, in which we observed a reduction in 3-linked mannopyrannosyl residues as a fraction of the total (Supplemental Table S3), indicating fewer unmodified backbone mannose residues.

## DISCUSSION

How *C. neoformans* interacts with host cells is a critical determinant of infection. In this study, we screened 4,692 cryptococcal deletion strains and identified 93 that demonstrated altered uptake by both mouse and human primary phagocytes. GO analysis showed that this set was enriched in sequences that encode proteins with roles in cytoskeletal interactions, protein transport, and organization of the cell wall and capsule. This is consistent with our understanding that the outer layers of the cell and secreted factors modulate interactions with host cells. We also identified strains with defects in only one host system. Why the species differ in this regard could be a fruitful area for future research.

Capsule is a critical factor in the interaction between *C. neoformans* and host cells, but the mechanisms of its synthesis and transport remain largely unknown. To approach this question, we performed a secondary screen for altered capsule thickness, which yielded 131 mutants. For comparison, a previous screen of 1,201 deletion strains, using colony morphology as the primary screen, yielded 16 capsule-related genes (19). Furthermore, our hits represent approximately one third of the original set of uptake mutants that we tested. This shows that our strategy of using these mutants as the starting population to screen for capsule defects was highly successful in enriching for capsule phenotypes. Notably, 42 of the hits are annotated as encoding hypothetical proteins (49), which may be subjects for future investigation. The others were enriched for GO terms related to cell wall and capsule organization, which validated our methods, and for terms involving transporter activity, intracellular transport, and phosphatidylinositol binding. As our results with Sac1 demonstrate, the study of such processes, which are not directly related to capsule production, may still yield insight into capsule biosynthesis.

Interestingly, over half of our capsule screen hits were hypercapsular strains. This highlights an important strength of our image-based technique, as prior methods (including colony morphology, antibody blotting, or volume-based methods) identify strains that lack capsule more readily than hypercapsular cells (50). That said, image-based screens may also have weaknesses. For example, mutants with general morphological defects may register as false positives, as cells that fail to divide might appear to be one large cell with a capsule defect. Further, this method cannot identify mutants that are completely acapsular, as they will not bind antibody.

Of the 45 gene products identified as hits in all three of our screens, the largest subset of the uncharacterized hits was predicted to be involved in the secretory pathway. We focused on one of these hits, Sac1, a PI4P phosphatase encoded by *CKF44_06080.* The *CNAG_06080* gene (which encodes Sac1 in the closely-related reference strain H99) was previously identified as a site where random integration of a drug marker yielded an attenuated *C. neoformans* strain (51). Like our KN99 mutant, this strain showed reduced virulence, although direct comparison is limited because the mutations, mouse models, and experimental designs differ. Other investigators had proposed that this gene was essential, because they were unable to delete it (52). We expect this difference from our findings was due to the challenge of working with cells that have significant defects, although it could also reflect different strain backgrounds (H99S vs KN99) (53, 54).

By expressing and localizing a fluorescent PI4P-binding peptide (41, 55, 56), we supported a role for Sac1 in PI4P turnover in *C. neoformans*. Although we did not test this activity directly, based on homology to the well-characterized *S. cerevisiae* Sac1 protein, conservation of Sac1 function throughout eukaryotes, and our binding data, we believe PI4P is the main substrate of Sac1 in *C. neoformans*. It is curious that despite its high amino acid similarity to the *S. cerevisiae* Sac1 protein, over-expression of the latter did not rescue the defects of *C. neoformans sac1*Δ cells; this result may warrant further experimentation.

When s*ac1* mutants are grown in host-like medium, they develop large structures which we termed SLBs. The interiors of SLBs stain with Nile Red, which binds neutral lipids, and the edges stain with the sterol-binding dye filipin. SLBS thus share certain features with lipid droplets, which are composed of a neutral lipid core of mainly triacylglycerols and sterol esters surrounded by a phospholipid monolayer (57). More specifically, SLBs resemble “supersized” lipid droplets, which form in model yeast strains defective in phospholipid metabolism (46, 58). This phenomenon has also been observed in *C. neoformans* following treatment with the antidepressant sertraline (59). However, our filipin staining suggests the presence of ergosterol or sterol intermediates (60) on the perimeter of the SLBs, which has not been reported for regular or supersized lipid droplets; accordingly we have not adopted this terminology.

Ergosterol is distributed in a steeply increasing gradient from the ER to the plasma membrane (61). One model proposes that PI4P turnover by Sac1 maintains this gradient (62). If this mechanism occurs in *C. neoformans,* loss of PI4P turnover would likely result in the inability to shuttle ergosterol up its concentration gradient with consequent accumulation in the ER. Another model suggests that the ergosterol gradient depends on membrane phospholipid composition (61). In either case, loss of Sac1 could perturb lipid transport and homeostasis and lead to mislocalization of ergosterol.

PI4P is a key signaling lipid, that serves as a pro-secretory marker and modulates the exit of secretory traffic from the *trans-*Golgi (27, 44, 63). We propose that the lack of PI4P turnover in *sac1* mutants leads to its accumulation in the early portions of the cryptococcal secretory system. This would be consistent with our observations of reduced protein secretion and mislocalization of the Golgi-resident protein Cxt1 in *sac1*Δ cells. The Golgi serves as a central hub for trafficking of both proteins and lipids, and in doing so plays a key role in membrane trafficking and lipid metabolism (64). We hypothesize that SLAB formation in *sac1* mutants represents the failure of lipid trafficking under stressful, nutrient-limited host-like conditions, leading to the accumulation of lipid cargo and/or lipid metabolic precursors. SLAB formation and the poor growth of *sac1*Δ cells is prevented by fatty acid supplementation, which somehow bypasses the defect caused by PI4P accumulation and allows the cells to grow normally. How this occurs will be an interesting question for future research. Importantly, while fatty acid supplementation rescues SLAB formation in *sac1*Δ cells, capsule production is not restored. This key finding shows that the adverse effects of Sac1 loss occur in divergent pathways.

Capsule material is exported in secretory vesicles (24), so defects in anterograde traffic would also impair this process. Beyond reduced GXM secretion, which is consistent with this expectation, we observed alterations in GXM composition: the mannose backbone had a large increase in glucuronic acid substitution and a slight increase in xylose. We hypothesize that altered localization of capsule biosynthetic enzymes changes their access to nascent capsule material and precursor molecules. It is not known whether capsule polysaccharides traverse the cell as subunits or extended polymers (65, 66), but in either case increased exposure to biosynthetic enzymes could result in compositional changes such as we observed. Both the reduction of capsule size and changes in composition in *sac1* mutants likely contribute to their increased uptake by host phagocytes.

In this study, we identified *C. neoformans* mutants impaired in host interactions with primary cells and assessed the hits for capsule defects. This strategy greatly increased the number of sequences known to influence capsule production, including many previously uncharacterized genes. Analysis of these genes and the corresponding mutants will help us understand the processes required for synthesis of this central virulence factor, as exemplified in this work on Sac1. Importantly, two-thirds of our original uptake hits did not have altered capsule thickness, implicating additional mechanisms in this fundamental host-pathogen interaction. Further studies will be required to determine what leads to the altered uptake of these strains, which may include changes to the cell wall or surface mannoproteins (29) or structural alterations in the capsule that do not impact its thickness.

## METHODS

### Strain construction, validation, and growth

*sac1Δ*, *SAC1* complement, and *SAC1^D456A^*strains were made with split-marker biolistic transformation (67) in KN99α. Cxt1 was tagged with mNEONGREEN (pBHM2404, Addgene #173441, a gift from Hiten Madhani) using CRISPR with short homology-directed repair (68). FAPP1-mNeonGreen (from pRS406-PHO5-GFP-hFAPP1_PH_domain, Addgene #58723, a gift from Tim Levine) and *S. cerevisiae SAC1* strains were similarly inserted in the Safe Haven 2 region (68, 69).

Transformants were validated by antifungal resistance, PCR and/or whole genome sequencing. Expression of *S. cerevisiae* S288C *SAC1* was confirmed by extracting total RNA with TRIzol Reagent, synthesizing cDNA using SuperScript III (Thermo Fisher, #18080400), and PCR-amplifying a unique region of the gene (Fig. S1B).

For all assays except screening, *C. neoformans* strains were grown on YPD plates (2 days, 30°C) and single colonies were inoculated into YPD and grown overnight (30°C, shaking). For capsule inductions, cells were washed in PBS, inoculated in DMEM at 10^6^ cells/mL, and grown for 16-24 hours (static, 37°C, 5% CO_2_). For screening, cells were cultured from glycerol stocks in 96-well plates, grown overnight in YPD, and then sub-cultured and grown 24 hours in YPD. Detailed methods, including stress media and lipid supplementation, are in Supplemental Materials.

### Screening

For uptake screening, C57BL/6J mouse BMDMs were harvested as in (70) and differentiated for 9 days before isolation. HMDMs were from anonymous donors. PBMCs were isolated by Ficoll-Paque centrifugation and differentiated for 6 days. The screen assay was modified from (23). Briefly, opsonized Lucifer Yellow-stained fungi were added to host cells at an MOI of 20 (BMDM) or 30 (HMDM) and the cultures were co-incubated for one hour and washed. Adherent cells were stained with Calcofluor White; wells were washed, fixed, and permeabilized; and host cell nuclei were stained with propidium iodide before imaging on a BioTek Cytation 3. The uptake index (internalized cells per macrophage) was corrected for the number of fungal cells added and then normalized to the plate median (first round) or KN99 value (later rounds).

For capsule screening, strains identified as hits in the first round BMDM uptake screen were assessed for capsule size differences using slight modifications of our previous method (30). For this, cells were induced as above, capsules were stained with anti-GXM mAb-302-AF488, and cell walls were stained with Calcofluor White. Images were collected as above, and capsule thickness measured as in (31).

Further details of methods and hit selection for both screens are in Supplemental Material.

### Virulence studies

All animal protocols were approved by the Washington University Institutional Animal Care and Use Committee (Protocol #20-0108) and care was taken to minimize animal handling and discomfort. 6-8-week-old female C57BL/6 mice (The Jackson Laboratory) were anesthetized by subcutaneous injection of 1.20 mg ketamine and 0.24 mg xylazine in 110 μL sterile PBS. For 14-day infections, mice were intranasally infected with 5 × 10^4^ cryptococcal cells. For time course infection, mice were infected with 1.25 × 10^4^ cryptococcal cells and sacrificed at 6, 12, and 18-days post infection. For both infections, the lungs and brains were harvested, homogenized, and plated on YPD agar to assess fungal organ burden.

### Assays

Acid phosphatase secretion was measured as in (24). 10^7^ cells/mL cultures were grown in MM-KCl (0.5% KCl, 15 mM glucose, 10 mM MgSO_4_⋅7H_2_O, 13 mM glycine, 3.0 µM thiamine). Supernatants were sampled, incubated with substrate buffer (2.5 mM para-nitrophenylphosphate in 50 mM acetate buffer, pH 4.0) at 37°C for 5 minutes, and stopped with 800 µL saturated Na_2_CO_3_. Acid phosphatase secretion was quantified as A_420_/OD_600._ GXM ELISAs were performed as in (48). Cultures were induced in DMEM,and the supernatants filtered (0.2 μm). Flat-bottom Immulon 1B plates (Thermo Scientific) were coated with 1 μg/ml anti-GXM antibodies 339 and F12D2γ, washed with PBS with 0.5% TWEEN 20 (PBST), and blocked for 90 min with 1% BSA in PBST. Filtrates were diluted in PBST and incubated on the plate for 90 min. The plate was washed three times, probed with HRP-labeled mAb 339/F12D2γ, incubated with KPL TMB Microwell Peroxidase Substrate (SeraCare), developed, and quantified using A_450_.

### Imaging

For capsule visualization, cells were induced as above, mixed with India ink in PBS (1:2, v:v), and imaged by light microscopy. Capsules of at least 96 cells from randomly-chosen fields were measured using ImageJ. Nile Red and filipin staining was performed as in (47), and viewed on a ZEISS Axio Imager M2 fluorescence microscope. FAPP1-mNEONGREEN and Cxt1-mNEONGREEN stains were imaged with a Zeiss LSM880 confocal microscope. Morphological classification of Cxt1-mNeonGreen and Nile Red staining was performed on at least 94 cells from randomly-selected fields. Electron microscopy was as in (47) with one change detailed in Supplement.

### GXM analysis

GXM was isolated using precipitation with hexadecyltrimethylammonium bromide as in (71). Isolated material was hydrolyzed with trifluoroacetic acid (3 hr, 100°C) and composition assessed on a Dionex ICS-6000 instrument. Linkage analysis was performed by the Complex Carbohydrate Research Center (method modified from (72)).

## ACKNOWLEDGEMENTS

We thank Nicole Howard and Suhas Bobba for assistance with host cells, Todd Fehniger and lab for LRS chambers for HMDM isolation, Yan Xie for statistical analysis, Wandy Beatty for electron microscopy, and Tim Levine for plasmid sequences. We appreciate the assistance of Jessica Tuck and Alex Caron with phenotypic characterization and microscopy studies. We thank Doering lab members for discussions, and particularly Liza Loza, Daphne Boodwa-Ko, Thomas Hurtaux, and Zanetta Chang for comments on the manuscript.

This work was supported by NIH grants AI140979, AI135012, and AI178330 to T.L.D. E.A.G and H.L.C were partially supported by T32 GM007067, and E.A.G. by a Sondra Schlesinger Graduate Fellowship from the Department of Molecular Microbiology, Washington University School of Medicine. The cryptococcal deletion library, developed by the Madhani lab with funding from R01AI100272, was purchased from the Fungal Genetics Stock Center. The CCRC is supported by the U.S. Department of Energy, Office of Science, Basic Energy Sciences, grant number DE-SC0015662 to Parastoo Azadi.

## SUPPLEMENTAL MATERIAL FOR GAYLORD ET AL

### Supplemental Methods

#### BMDM preparation

BMDMs were isolated as in (70) from 9-week old C57BL/6J mice and frozen in 90% FBS, 10% DMSO at 10^7^ cells/mL. Thawed cells were differentiated for 9 days in RPMI-1640 with L-glutamine (Sigma) with 30% L-cell supernatant and 20% heat-inactivated fetal bovine serum (Gibco), with regular media changes. BMDM were resuspended in 10 mL ice cold PBS and isolated using 2.5 μg anti-mouse F4/80 biotin (clone BM8, eBiosciences) plus 0.5 μg mouse BD FC-Block (BD Pharmingen) with anti-biotin microbeads (Miltenyi Biotec) and a MACS separation column (Miltenyi Biotec).

#### Human macrophage preparation

Leukoreduction system (LRS) chambers were obtained from anonymous donors and human peripheral blood mononuclear cells (PMBC) were isolated by Ficoll-Paque centrifugation (Lymphoprep; Axis-Shield). 5×10^7^ PMBC were allowed to adhere to a treated culture dish (Corning) for 2 hours at 37°C with 5% CO_2_. Detached cells were removed, and adherent cells were maintained in RPMI-1640 differentiation media containing L-glutamine (Sigma), 10% fetal bovine serum (Sigma), 1% penicillin streptomycin (Fisher), 1% sodium pyruvate (Corning), 50ng/mL human recombinant M-CSF (Fisher), and 1% MEM non-essential amino acids (Corning). Medium was replaced every two days and adherent cells were harvested on day 6 using Cell Stripper (Corning).

#### Uptake screen

Assays were performed using a modification of our prior method (23). Host cells were seeded in flat clear bottom black polystyrene TC-treated 96-well plates (Corning) at a density of 10^5^ cells/well and incubated for one day at 37°C with 5% CO_2_. *C. neoformans* strains were cultured from frozen stocks in glycerol/Yeast Extract-Peptone-Dextrose medium (YPD) in 96 well format for 18-24 h at 30°C with shaking (800 rpm) (Southwest Science, SBT1500-H). At least 2 μL of cells were then sub-cultured in 100 μL YPD and grown for 18-24 h as above. After sub-culture, cells were sedimented (3000 g, 2 min, RT), washed twice with 200 μL of PBS, washed once with 200 μL McIlvaine’s Buffer [pH 6], and resuspended in 80 μL of the same. Cells were then stained with 5 mg/mL Lucifer Yellow (30 min, RT, 650 rpm) and resuspended in 100 μL of PBS. To account for variation in growth rates of *C. neoformans* mutants, OD_600_ was used to roughly standardize cell number and maintain results in the linear range of the assay. To do this, individual mutants were categorized as having high, medium, or low growth based on the OD_600_ after resuspension in PBS and the average value of each group was used to prepare wells with approximately 10^7^ fungal cells/mL in PBS. Fungal cells were then opsonized with 10% by volume C57Bl6/J mouse serum (for BMDM) or human serum from male AB plasma (for HMDM, Sigma) for 30 minutes at 37°C at 250 rpm. Opsonized cells were then added to wells containing RPMI and macrophages at a multiplicity of infection of 20 (BMDM) or 30 (HMDM). Cells were co-incubated for one hour at 37°C with 5% CO_2_ to allow uptake and washed. This and the following washes were performed using a Biotek ELx405 Microplate Washer (2 washes, PBS). Adherent fungal cells were stained with 150 μL of 5 μg/mL Calcofluor White (Sigma) (15 min, 4°C, static). Samples were then washed, fixed with 4% formaldehyde (10 min, RT), washed, permeabilized with 1 mg/mL saponin (20 min, RT), washed, and stained with 50 μg/mL propidium iodide (15 min, RT). Imaging was performed on a BioTek Cytation 3 imager collecting 12 images per well in a 4×3 grid pattern.

A separate plate of each *C. neoformans* strain alone was counted to allow normalization for the number of cells of each mutant strain that were used in the assay. The phagocytic index for each mutant was determined by calculating the number of internalized cells per macrophage and normalizing this to the value for WT KN99 and the number of fungi added. An uptake score was then calculated for each well as fold-change of the normalized phagocytic index compared either to the plate median (for the initial screen of Madhani library plates) or to WT (for the follow-up screens with BMDM and HMDM). For the initial screen of the Madhani deletion collection, we calculated the first quartile (Q1), third quartile (Q3), and interquartile range (IQR) for each plate. We defined hits as those below Q1-c*IQR or above Q3-c*IQR, where c was calculated using a targeted error rate, α, of 0.01. For the second round of screening in BMDM and HMDM, we used an internal WT control and defined hits as those strains whose uptake scores differed from WT in a statistically significant way, using Dunnett’s 1-way ANOVA and a p value cutoff of 0.05.

#### Capsule screen

Strains identified as hits in the uptake assay and wild type control cells were cultured in YPD from frozen stocks in glycerol/YPD in 96 well format (ON, 30 °C, 800 rpm); 10 μL aliquots were then sub-cultured into 190 μL fresh YPD and grown for an additional 24 h. Cells were then sedimented (3000 g, RT) and resuspended in 200 μL DMEM. To achieve the low cell density necessary for optimum capsule induction, the average OD_600_ of the plate was used to calculate the volume of cells to resuspend in 150 μL DMEM to achieve an average OD_600_ of 0.01. Capsule induction was performed in a 96-well glass-bottom plate coated with poly-lysine (Eppendorf) and incubated for 24 h at 37°C with 5% CO_2_. KN99α cells grown overnight in YPD were added to empty wells 10 min before the induction ended to serve as an uninduced control. Following induction, cells were stained and imaged as published previously (30). Briefly, the capsules were stained with anticapsular monoclonal antibody 302 conjugated to Alexa Fluor 488 (Molecular Probes) and the cell walls stained with Calcofluor White (MP Biomedicals). Imaging was performed on a BioTek Cytation 3 imager collecting 64 images per well in an 8×8 grid pattern. Images were prepared, annotated, and cell wall and capsule diameters for annotated cells calculated as described previously (30). Capsule thickness was defined as the difference between cell wall and capsule diameters. A negative control value (mean capsule thickness of WT cells grown in YPD in the same plate) was subtracted and the result normalized to the value for similarly-corrected positive control samples (WT cells induced in the same plate) to yield a capsule score. Hypocapsular and hypercapsular hits were categorized as those with capsule scores of less than 0.5 or more than 2.0, respectively.

#### Gene ontology enrichment analysis

Gene ontology (GO) analysis was performed using FungiDB, using both manually curated and computationally predicted GO terms (49, 73, 74). GO categories that were enriched with a P-value cutoff of 0.01 were manually selected for each screen to minimize over-representation of redundant phenotypes.

#### Strain construction and validation

*sac1Δ*, *SAC1* complement, and *SAC1^D456A^* strains were engineered using biolistic transformation with a split-marker strategy (67) in a KN99α background. Design of the catalytically-inactivated construct was based on alignment (75) to the *S. cerevisiae* S288C Sac1 amino acid sequence. mNeonGreen-tagged strains were generated using pBHM2404, a gift from Hiten Madhani (Addgene 173441). PI4P-binding constructs were generated using pRS406-PHO5-GFP-hFAPP1 PH domain, a gift from Tim Levine (Addgene 58723) (41). This sequence was placed between a *TEF1* promoter and terminator and inserted into the Safe Haven 2 (SH2) region of the genome (69). All fluorescent constructs were generated using CRISPR with short homology-directed repair (68). Transformants were confirmed by appropriate antifungal resistance, PCR, and whole genome sequencing.

To test complementation of *sac1*Δ by the corresponding *S. cerevisiae* sequence, the *SAC1* gene was amplified from *S. cerevisiae* strain S288C and inserted into plasmid pMSC042-neo between an *ACT1* promoter and *TRP1* terminator. This construct (containing the promoter, *SAC1* sequence, terminator and antifungal resistance marker) was inserted into the SH2 region of WT and *sac1*Δ cells using CRISPR (68, 69). Total RNA was extracted from cells grown in YPD and DMEM using TRIzol Reagent (Thermo Fisher) and cDNA was synthesized using the SuperScript III First-Strand Synthesis SuperMix kit (Thermo Fisher). PCR amplification of an S288C Sac1-specific sequence from the cDNA confirmed expression of the S288C *SAC1* gene in both backgrounds (Fig. S1B).

#### Fungal cell growth and capsule induction

*C. neoformans* strains were grown from glycerol stocks on YPD plates for 2 days at 30°C. Overnight cultures were inoculated from single colonies into 4 mL of YPD medium and grown overnight at 30°C with shaking (unless otherwise noted). For all capsule induction experiments, overnight cultures were washed in sterile PBS, inoculated into DMEM at 10^6^ cells/mL, and grown statically at 37°C with 5% CO_2_ for 16-24 hours (as indicated in the text).

#### Virulence studies

All animal protocols were approved by the Washington University Institutional Animal Care and Use Committee (Protocol #20-0108) and care was taken to minimize animal handling and discomfort. 6-8-week old female C57BL/6 mice (The Jackson Laboratory) were anesthetized by subcutaneous injection of 1.20 mg ketamine and 0.24 mg xylazine in 110 μL sterile PBS. For 14-day infections, mice were intranasally infected with 5 × 10^4^ cryptococcal cells. For time course infection, mice were infected with 1.25 × 10^4^ cryptococcal cells and sacrificed at 6-, 12-, and 18-days post infection. For both infections, the lungs and brains were harvested, homogenized, and plated on YPD agar for calculation of fungal organ burden.

#### Stress plate dot spotting

Overnight cultures were pelleted and washed twice in PBS, then diluted to 10^7^ cells/mL. 4 μL of this stock and four serial 1:10 dilutions were spotted onto YPD agar and incubated at both 30 and 37°C for 48 hours. To impose plasma membrane stress, the YPD was supplemented with 0.01% sodium dodecyl sulfate or 1 µg/ml amphotericin B; to impose cell wall stress, the YPD was supplemented with 0.05% Congo red.

#### Acid phosphatase secretion assay

Single colonies were inoculated into 5 mL YPD, grown shaking at 30°C for 6 h, sub-cultured into 25 mL MM-KH_2_PO_4_ (0.5% KH_2_PO_4_, 15 mM glucose, 10 mM MgSO_4_⋅7H_2_O, 13 mM glycine, 3.0 µM thiamine), and grown overnight at 30°C. Cells were then pelleted, washed twice with MM-KCl (0.5% KCl, 15 mM glucose, 10 mM MgSO_4_⋅7H_2_O, 13 mM glycine, 3.0 µM thiamine), and resuspended in 50 mL MM-KCl at 10^7^ cells/mL. Cultures were grown at 30°C with shaking, and 2 mL aliquots were collected at each timepoint. For each, pelleted cells were resuspended in 400 µL substrate buffer (2.5 mM paranitrophenylphosphate in 50 mM acetate buffer, pH 4.0, Sigma 71768-5G), incubated at 37°C for 5 minutes, and the reaction stopped with 800 µL saturated Na_2_CO_3_. Acid phosphatase secretion was quantified using absorbance at 420 nm, corrected by subtraction of a cell-free blank value, and normalized to cell density using OD_600_.

#### GXM ELISA

Cultures were induced in DMEM for the indicated times, at which point cells were removed by centrifugation (3000 x g, 5 min, RT), the supernatant fraction filtered through a 0.2μm filter, and the filtrate diluted 1:500 in PBS. Flatbottom Immulon 1B microtiter plates (Thermo Scientific) were coated overnight with a 1 μg/ml mix of anti-GXM antibodies 339 and F12D2γ in PBS. This and the following incubations were performed at room temperature. The plates were washed twice with 200 μL PBS with 0.5% TWEEN 20 (PBST), blocked for 90 min with 200 μL 1% BSA in PBST. The blocking buffer was replaced with 100 μL PBST and 100 μL of filtered sample was added and incubated for 90 min. The plate was then washed three times as above and the bound GXM was probed with HRP-labeled mAb 339/F12D2γ, incubated for 1 minute with KPL TMB Microwell Peroxidase Substrate (SeraCare), developed, and quantified using absorbance at 450 nm.

#### Imaging

For fluorescence and confocal microscopy, cells were grown as indicated above and washed twice in PBS. For capsule visualization, cells were resuspended in India ink in PBS (1:2, v:v) and imaged on a ZEISS Axio Imager M2 fluorescence microscope. Capsule thickness was measured manually in ImageJ, using a total of at least 96 cells from at least six randomly-chosen fields of view. To stain neutral lipids, a 0.01% Nile Red stock in acetone was diluted to a working concentration of 0.0005% in PBS. Cells were incubated in 1 mL working Nile Red solution for 15 min in the dark, rotating. Cells were washed twice in PBS, resuspended in PBS, and imaged as above. For quantification of morphology, a minimum of 60 cells from randomly-chosen slide coordinates were examined. For filipin staining, cells were incubated in a freshly prepared solution of 5 μg/mL filipin (Millipore Sigma, F9765-25MG) in DMSO for 15 minutes at room temperature, rotating, then washed twice with PBS before imaging as above. To visualize FAPP1-mNeonGreen and Cxt1m-NeonGreen, cells were imaged using a Zeiss LSM880 confocal microscope. For quantification of morphology, a minimum of 94 cells from randomly-chosen slide coordinates were examined. For electron microscopy, cells were grown overnight in YPD or induced for 24 hours in DMEM before processing as in (47), except that the first incubation in 0.1 M sodium cacodylate buffer was performed for 2 h at RT.

#### Lipid supplementation

For lipid supplementation, a fatty acid stock solution (10 mM in ethanol) was diluted 1:10 into 4 mg/ml BSA in DMEM and then sonicated (Bransonic 2 Ultrasonic Bath Sonicator) before addition to cultures as indicated. After 24-hour of incubation (37°C, 5% CO_2_), cells were serially diluted 1:10 and plated onto YPD agar, with the top spot representing the undiluted culture.

#### GXM purification and analysis

GXM was isolated using selective precipitation of culture supernatants with hexadecyltrimethylammonium bromide (CTAB) as described previously (48). Isolated material was hydrolyzed in trifluoroacetic acid for 3 hours at 100°C and compositional analysis performed using high-performance anion-exchange chromatography with pulsed amperometric detection on a Dionex ICS-6000 instrument (Thermo Scientific). Glycosyl linkage analysis was performed by the Complex Carbohydrate Research Center at the University of Georgia using combined gas chromatography/mass spectrometry of partially methylated alditol acetate derivatives following carboxyl reduction to detect uronic acids (modified from (72)).

## Supplemental Table Legends

**Supplemental Table S1.** Uptake screen results. (S1A) All results from the initial BMDM uptake screen. (S1B) Hits from the initial BMDM uptake screen. (S1C) All results from the second BMDM uptake screen. (S1D) Hits identified from the second BMDM uptake screen. (S1E) All results from the HMDM uptake screen. (S1F) Hits from the HMDM uptake screen.

**Supplemental Table S2.** Capsule screen results. (S2A) All results from the capsule screen. (S2B) Hits identified from the capsule screen. (S2C) Overlap hits identified in all three screens: HMDM, BMDM, and capsule.

**Supplemental Table S3.** Linkage analysis of independently isolated WT, sac1Δ, and *SAC1* strains. Values are percent area of relevant residue peaks. Numbers in header row refer to independently isolated samples.

| Residue | WT #1 | *sac1*Δ #1 | WT #2 | *sac1*Δ #2 | *SAC1* #2 |
| --- | --- | --- | --- | --- | --- |
| Terminal Xylopyranosyl residue (t-Xyl) | 12.0 | 19.7 | 10.4 | 16.2 | 11.1 |
| Terminal Mannopyranosyl residue (t-Man) | 1.3 | 0.3 | 0.3 | 0.3 | 0.4 |
| Terminal Glucopyranosyl uronic acid residue (t-GlcA) | 17.9 | 15.4 | 19.1 | 16.2 | 18.2 |
| 3-linked Mannopyranosyl residue (3-Man) | 29.9 | 23.3 | 27.8 | 21.1 | 30.6 |
| 2,3-linked Mannopyranosyl residue (2,3-Man) | 36.0 | 38.5 | 37.5 | 42.8 | 34.9 |
| 2,3,4-linked Mannopyranosyl residue (2,3,4-Man) | 2.9 | 2.8 | 4.9 | 3.5 | 4.8 |

## Supplemental Figure Legends

**Supplemental Figure S1.**
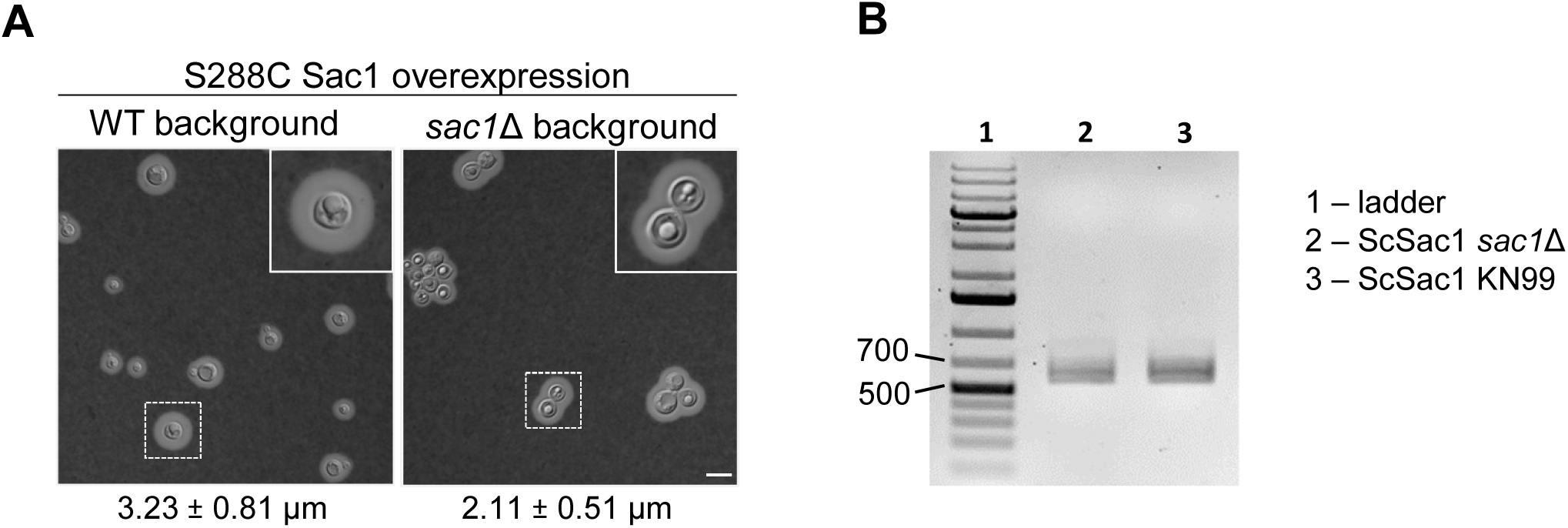
Overexpression of the *S. cerevisiae* Sac1 protein in *sac1*Δ does not restore *C. neoformans* capsule synthesis. **(A)** Negative stain of the indicated strains grown for 24 hours in DMEM at 37°C with 5% CO_2_, with the capsule thickness in μm (mean ± SD). Scale bar, 10 μm. **(B)** Amplification of a unique 651 bp region of the S288C Sac1 coding sequence from cDNA generated from fungal cells of the indicated strain grown in YPD shows that the *S. cerevisiae SAC1* gene is expressed in both *C. neoformans* strains.

**Supplemental Figure S2.**
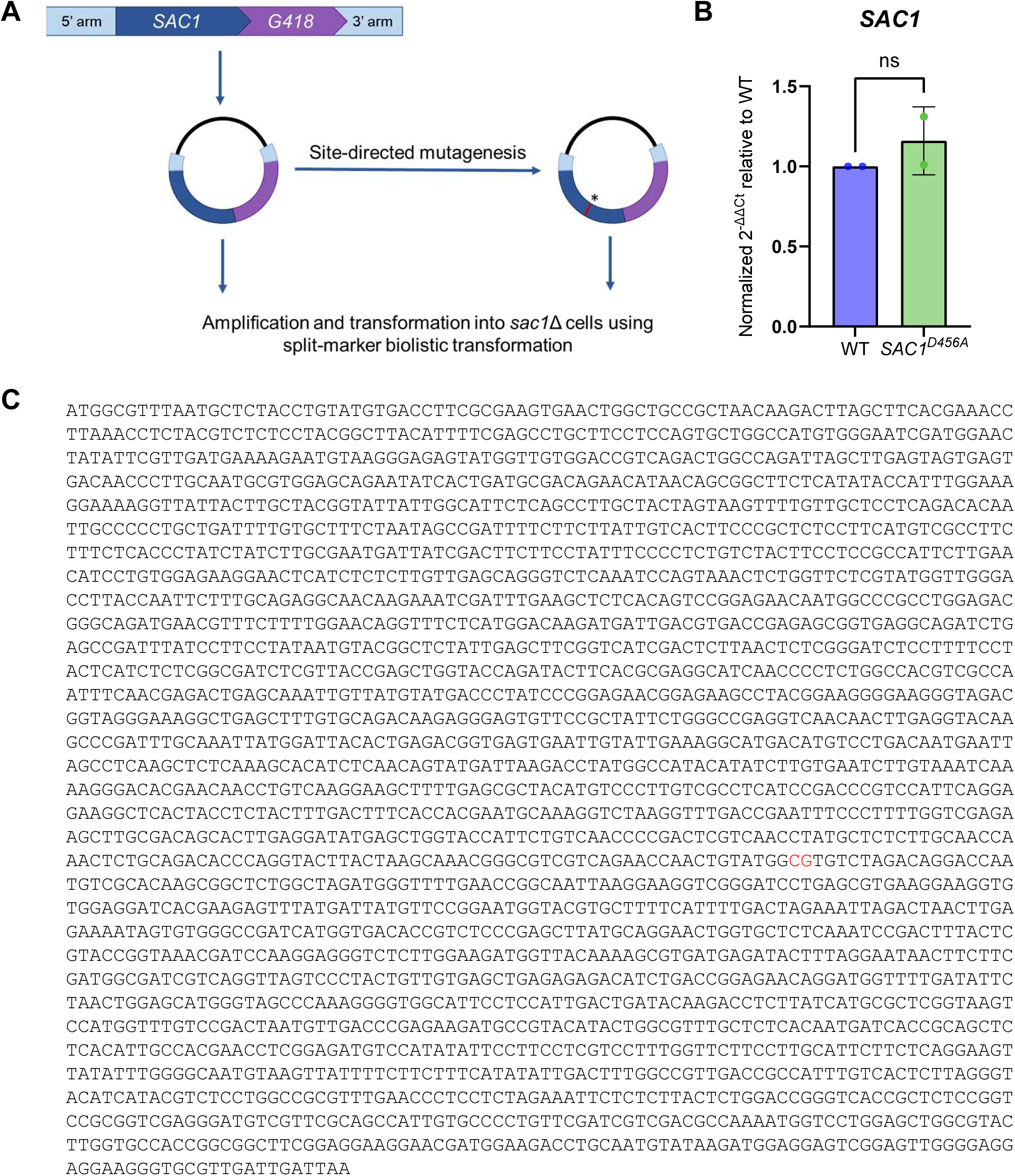
Construction and validation of the *SAC1D^456A^* strain. **(A)** Strain construction strategy for *SAC1* complement and *SAC^D456A^* inactivated complement strains. **(B)** Normalized expression of *SAC1* in WT and *SAC1^D456A^* cells grown for 24 hours in DMEM at 37°C with 5% CO_2_. **(C)** Nucleotide sequence of *SAC1^D456A^.* Red, nucleotide differences from WT *SAC1*.

**Supplemental Figure S3.**
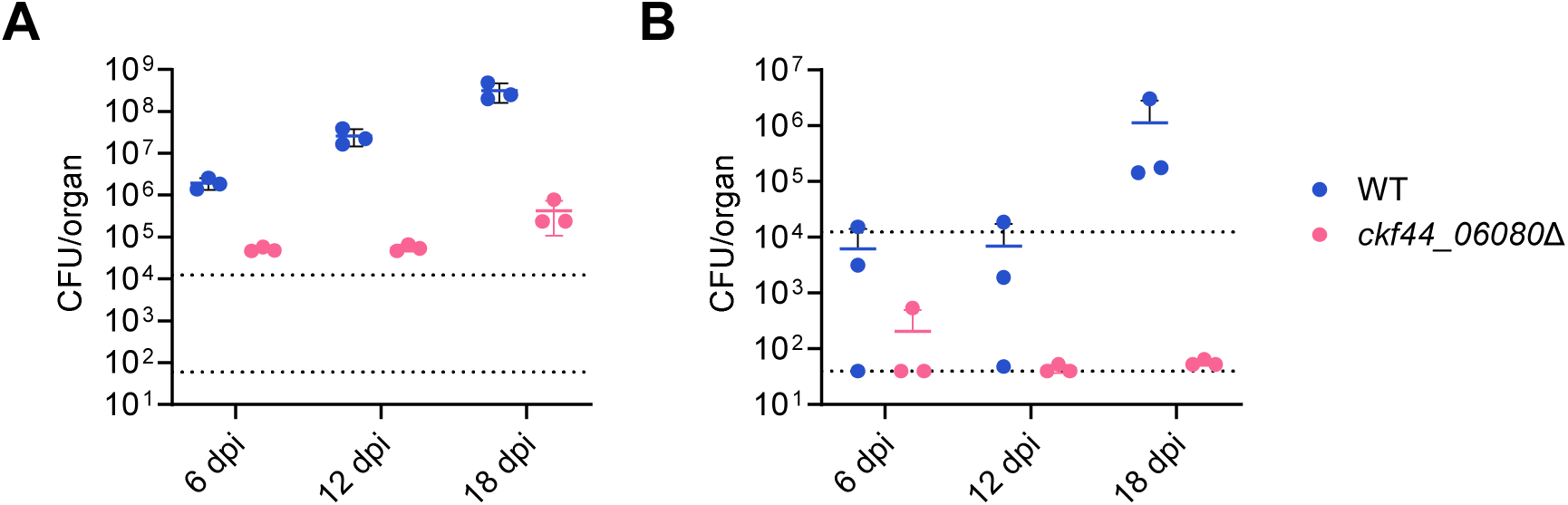
Cells lacking functional Sac1 have reduced virulence. **(A)** Lung and **(B)** brain burden of C57BL/6 mice sacrificed at the indicated days post infection (dpi) with 1.25×10^4^ fungal cells of *ckf44_06080*Δ from the Madhani *C. neoformans* deletion collection. Upper dotted line, inoculum. Lower dotted line, limit of detection. Each symbol represents one mouse.

**Supplemental Figure S4.**
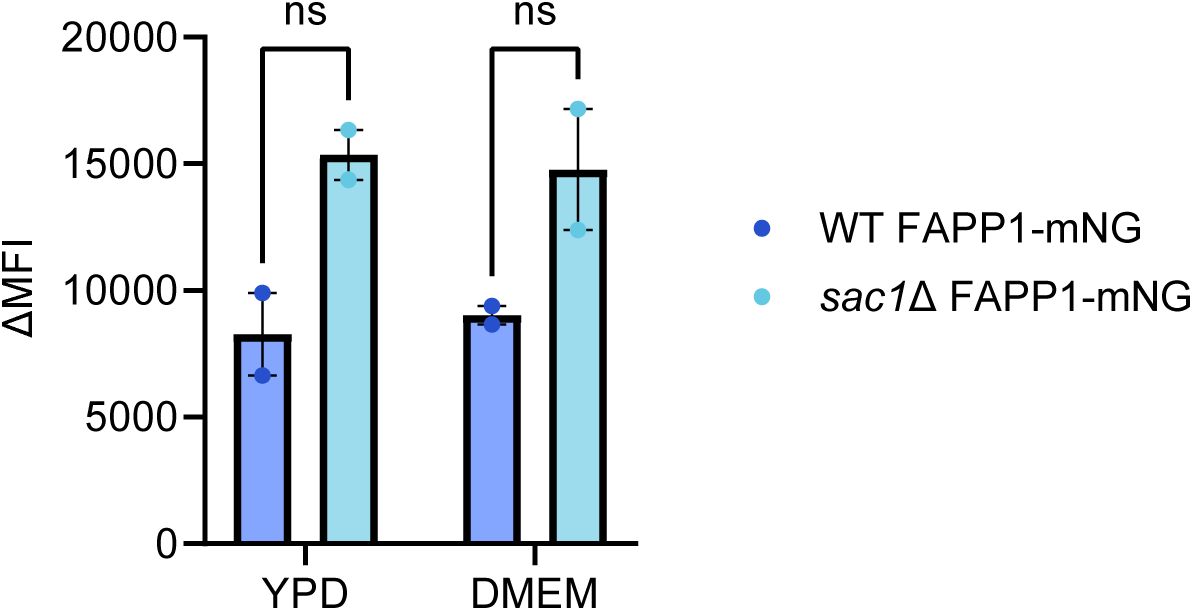
PI4P content of WT and *sac1*Δ strains. Difference in median fluorescence intensity between FAPP1-mNG expressing and control cells (ΔMFI) in WT and mutant backgrounds, grown in the indicated medium. ns, not significant by unpaired t test.

**Supplemental Figure S5.**
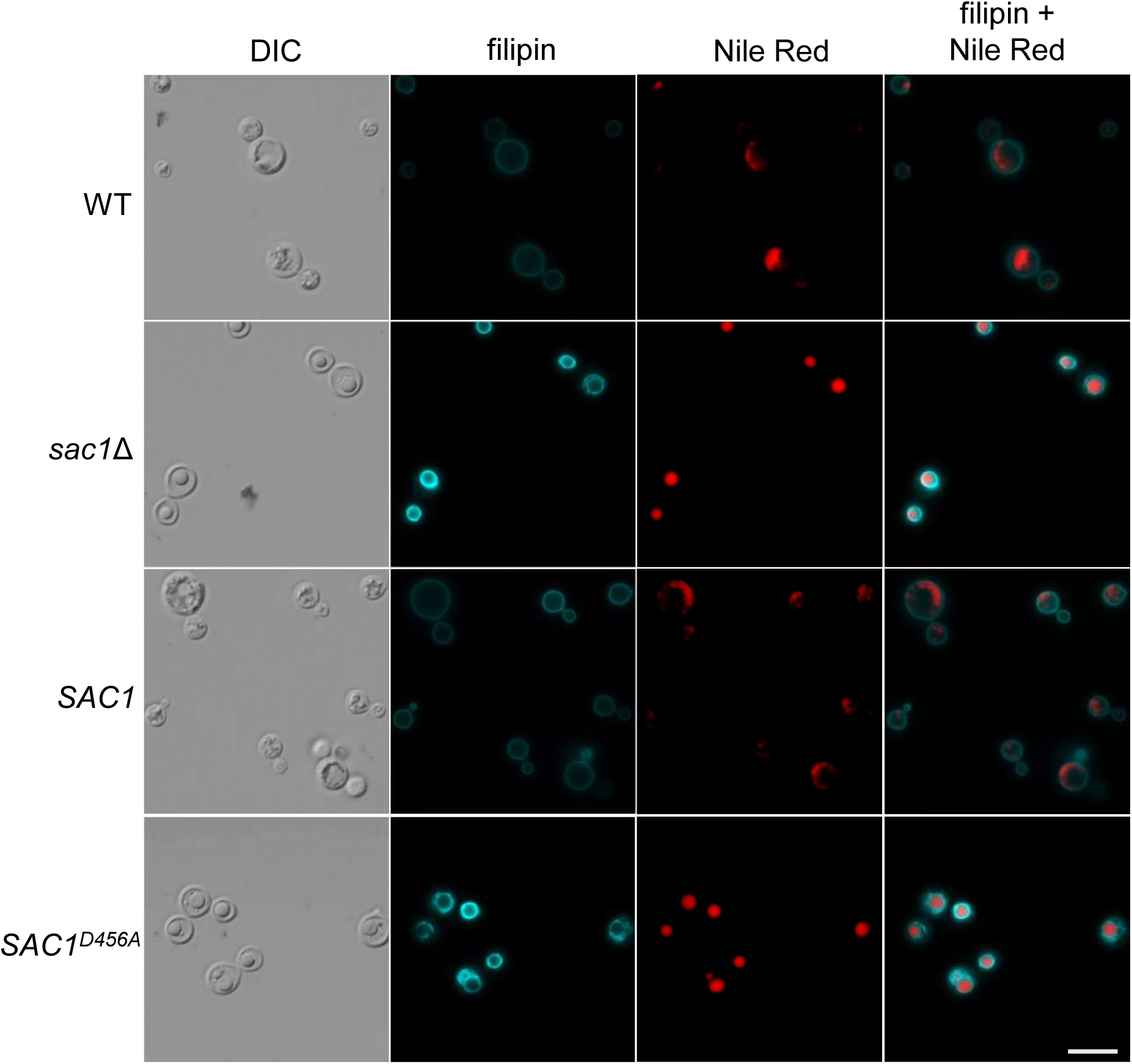
Dual staining with filipin and Nile Red. All cells were grown in DMEM for 24 hours at 37°C with 5% CO_2_. DIC, Differential interference contrast. Scale bar, 10 μm.

**Supplemental Figure S6.**
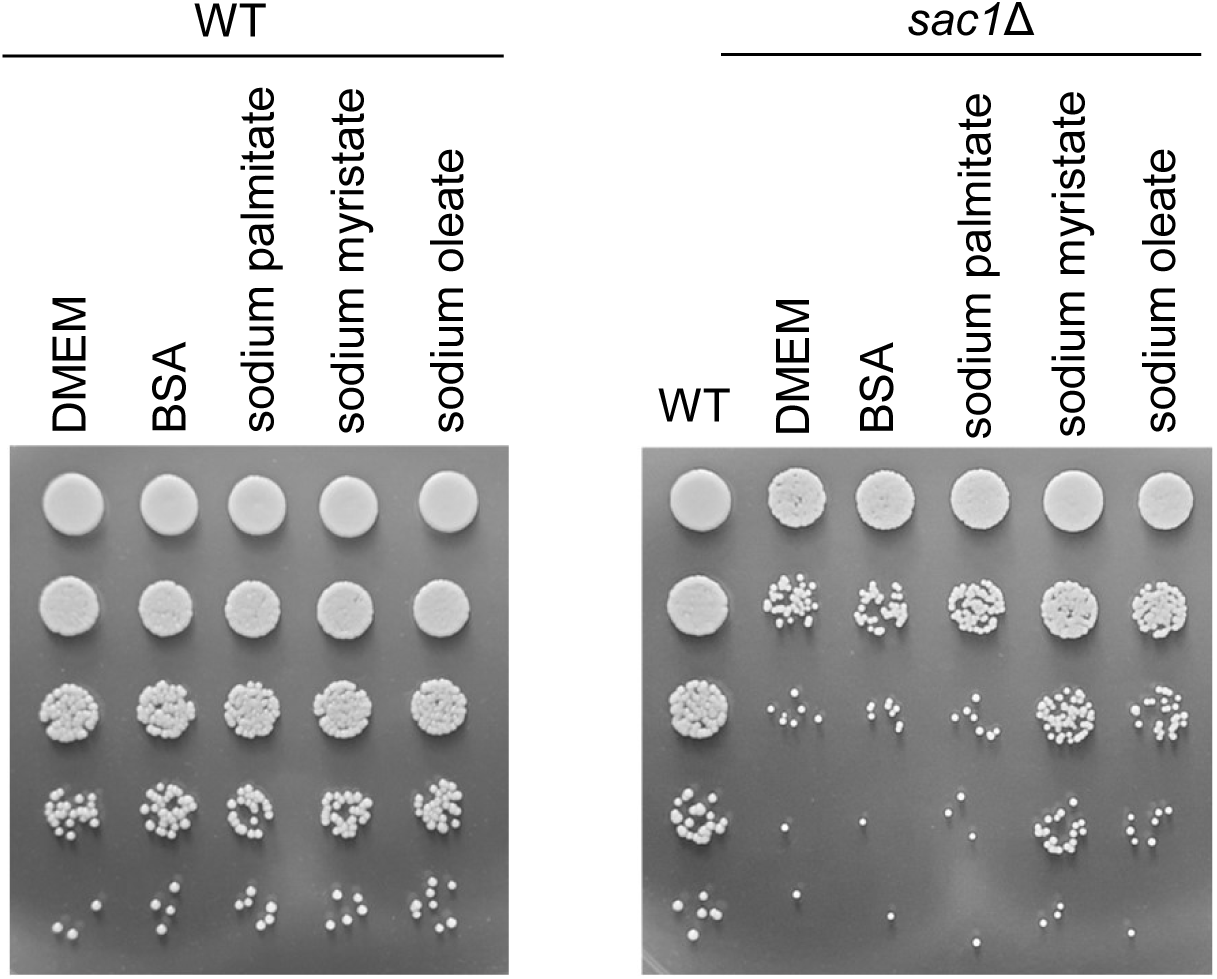
Supplementation with exogenous fatty acids partially restores growth of *sac1*Δ cells. Serial dilutions of (left) WT or (right) *sac1*Δ cells grown for 24 hours in DMEM alone, DMEM supplemented with BSA, or DMEM with 100 μM of the indicated fatty acid sodium salt conjugated to 0.4 mg/mL BSA.

## Notes

### Competing Interest Statement

The authors have declared no competing interest.

### Summary of Updates

Figures 1, 2, and 3 revised and/or updated; addition of Supplemental Figures 2 and 4; revision to Supplemental Table 2; text revisions.

